# Fully human single-domain antibodies against SARS-CoV-2

**DOI:** 10.1101/2020.03.30.015990

**Authors:** Yanling Wu, Cheng Li, Shuai Xia, Xiaolong Tian, Zhi Wang, Yu Kong, Chenjian Gu, Rong Zhang, Chao Tu, Youhua Xie, Lu Lu, Shibo Jiang, Tianlei Ying

## Abstract

The COVID-19 pandemic is spreading rapidly, highlighting the urgent need for an efficient approach to rapidly develop therapeutics and prophylactics against SARS-CoV-2. We describe here the development of a phage-displayed single-domain antibody library by grafting naïve CDRs into framework regions of an identified human germline IGHV allele. This enabled the isolation of high-affinity single-domain antibodies of fully human origin. The panning using SARS-CoV-2 RBD and S1 as antigens resulted in the identification of antibodies targeting five types of neutralizing or non-neutralizing epitopes on SARS-CoV-2 RBD. These fully human single-domain antibodies bound specifically to SARS-CoV-2 RBD with subnanomolar to low nanomolar affinities. Some of them were found to potently neutralize pseudotyped and live virus, and therefore may represent promising candidates for prophylaxis and therapy of COVID-19. This study also reports unique immunogenic profile of SARS-CoV-2 RBD compared to that of SARS-CoV and MERS-CoV, which may have important implications for the development of effective vaccines against SARS-CoV-2.

---

Recently, an outbreak of novel coronavirus (SARS-CoV-2) has spread rapidly around the globe ^1-4^. As of 29 March, 2020, there have been 634,835 laboratory-confirmed human infections globally, including 29,891 deaths (https://www.who.int/emergencies/diseases/novel-coronavirus-2019/situation-reports). This marks the third major outbreak caused by a new coronavirus in the past two decades, following severe acute respiratory syndrome coronavirus (SARS-CoV) and Middle East respiratory syndrome coronavirus (MERS-CoV). Furthermore, SARS-CoV-2 is one of the most transmissible coronaviruses identified so far, with the coronavirus disease (COVID-19) quickly accelerating into a global pandemic. These facts highlight the urgent need for an efficient approach to rapidly develop therapeutics and prophylactics against SARS-CoV-2, which could not only be potentially implemented in dealing with COVID-19 during the current outbreak, but also strengthen our preparedness and response capacity against emerging coronaviruses in the future.

Monoclonal antibodies (mAbs) are showing unprecedented value, and represent the largest and fastest-growing sector in pharmaceutical industry. During the previous SARS and MERS outbreaks, a number of neutralizing mAbs have been developed and proved their therapeutic potential in the treatment of coronavirus infections ^5-10^. Despite this, their clinical usefulness has been hampered by the time-consuming and costly antibody manufacturing processes in eukaryotic systems. The large-scale production of mAbs typically takes at least 3 to 6 months, making them difficult to be timely produced and used in an epidemic setting. An attractive alternative for mAbs is single-domain antibodies from camelid immunoglobulins, termed VHH or nanobodies that are the smallest naturally occurring antigen-binding protein domains with a molecular weight of 12-15 kDa ^11^. Their small size provides several advantages over conventional mAbs (150 kDa), including larger number of accessible epitopes, relatively low production costs, and easiness of rapid production at kilogram scale in prokaryotic expression systems. More importantly, nanobodies can be administered by inhaled delivery due to the small size and favorable biophysical characteristics, and thus are considered to be particularly suitable for the treatment of respiratory diseases ^12^. For instance, ALX-0171, an inhaled anti-respiratory syncytial virus (RSV) nanobody developed by Ablynx, was found to have robust antiviral effects and reduce signs and symptoms of RSV infection in animal models, and well tolerated at all doses when administered by inhalation in clinical trials ^13^. These findings confirmed the feasibility of administering nanobodies via inhalation. However, the camelid origin of nanobodies limits their application as therapeutics in human. To reduce the risk of immunogenicity, strategies for humanization of camelid nanobodies have become available in recent years but suffered from time- and labor-consuming ^11^. Besides, humanized nanobodies still retain a small number of camelid residues, especially those within the framework region 2 (FR2), in order to maintain the solubility and antigen-binding affinity of parental antibodies ^11,14^.

In contrast to the camelid nanobodies which are naturally devoid of light chains, heavy chain variable domains (VH) of conventional antibodies are paired with light chain variable domains (VL), and generally poorly expressed or easy to aggregate in the absence of light chains. It was proposed that several specific “hallmark” residues (F37, E44, R45, and G47) within FR2 may contribute to the high solubility and stability of isolated nanobodies ^15^. Interestingly, the analysis of 2391 nanobody sequences from a public database revealed that their FR2 regions are relatively divergent including the hallmark residues which have been considered to be strictly conserved (Fig. 1a). Furthermore, we and others have previously identified some isolated human VH single domains, which were independently folded and exhibited very similar biophysical properties to camelid nanobodies ^16,17^. These findings inspired us to revisit the structural feature of single-domain antibodies, and hypothesize that certain VH framework regions could compensate for the absence of VL, resulting in the soluble human single-domain antibodies. Therefore, we first searched the IMGT database for the human IGHV alleles sharing the same germline framework regions (FR1, FR2, or FR3) with m36, an HIV-1 neutralizing VH that was found to be highly soluble and stable. As a result, 17 human germline IGHV alleles, along with a camelid nanobody (VHH#3) as control ^18^, were cloned, expressed in *Escherichia coli*, and characterized for their biophysical properties (Fig. 1b). Nine out of 17 alleles could be highly expressed with yields of over 10 mg/L bacterial culture, and 10 out of 17 possess protein A binding capabilities. Notably, germline 3-66*01 exhibited the most advantageous properties, including comparable midpoint transition temperature (T_m_) to that of nanobody measured by intrinsic protein fluorescence, and the highest aggregation temperature (T_agg_) among all tested single-domain antibodies measured by static light scattering. These results confirmed the feasibility of using human single-domain antibodies as ideal alternatives to camelid nanobodies in therapeutic applications.

**Figure 1.**
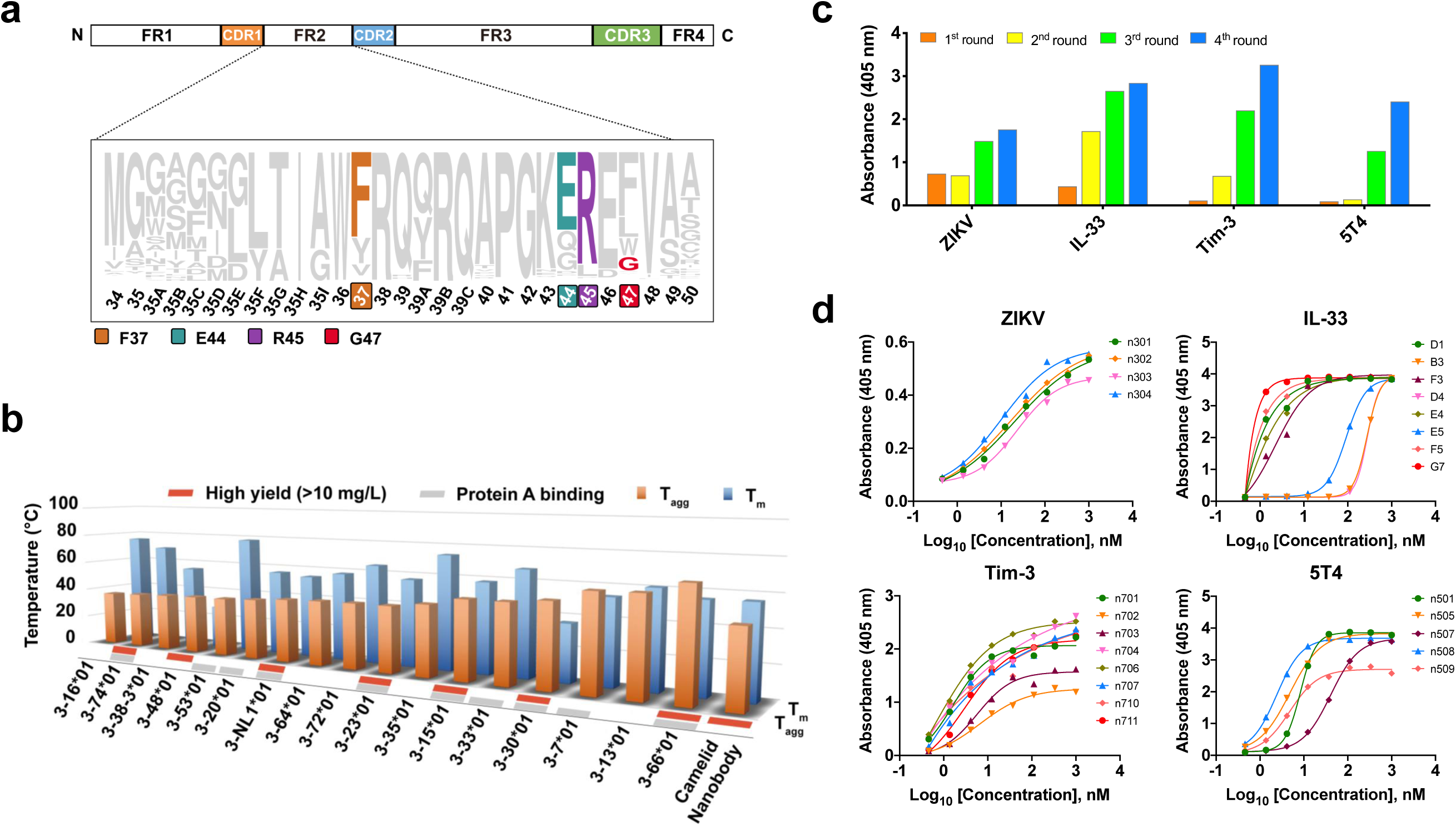
Development of a versatile platform for rapid isolation of fully-human single-domain antibodies. **a**, Representation of camelid nanobody framework (FR) and complementarity-determining (CDR) regions, showing the four hydrophilic amino acids (Phe37, Glu44, Arg45, Gly47) in the FR2 region that may contribute to high solubility and stability of isolated nanobodies. **b**, Characterization of biophysical properties (protein yield, protein A binding capacity, stability and aggregation) of 17 isolated human germline IGHV alleles along with a camelid nanobody. **c**, Polyclonal phage ELISA showing the binding of the first to fourth rounds of phages to target antigens. Bound phages were detected with anti-M13-HRP conjugate. **d**, Binding activity of purified single-domain antibodies against target antigens evaluated by ELISA.

Next, we aimed to establish a generalizable platform for rapid development of human single-domain antibodies. We used germline 3-66*01 framework regions as the scaffold for grafting of heavy chain CDRs cloned from several naïve antibody libraries. These libraries were previously constructed from the blood of healthy adult donors, and their effectiveness has been proved by the successful isolation of potent germline-like human monoclonal antibodies against various targets such as H7N9 avian influenza virus ^19^, MERS-CoV ^20^, and Zika virus ^21^. Consequently, such CDR grafting resulted in a very large and highly diverse phage-displayed single-domain antibody library (size ∼2×10^11^). To validate the quality of the library, several parallel bio-panning were performed against a set of representative antigens, including viral antigen, cytokine, and surface antigens on immune or tumor cells. In all the tests, potent phage enrichments were observed after two or three rounds of panning (Fig. 1c), and panels of single-domain antibodies could be identified with binding affinities in the low nanomolar/subnanomolar range (Fig. 1d). These antibodies are monomeric and could be solubly expressed at high levels in *Escherichia coli* with yields ranging from 15 to 65 mg/L culture. Moreover, their sequences are of fully human origin with minimal divergence from the germline predecessors.

This technology enabled us to rapidly develop fully human single-domain antibodies against SARS-CoV-2. To this end, the receptor binding domain (RBD) of SARS-CoV-2 was produced and biotinylated at a specific site for use as the target antigen during bio-panning. Significant enrichment was achieved after two rounds of panning, and a panel of 37 unique single-domain antibodies was identified using the soluble expression-based monoclonal ELISA. According to the sequence similarities among these antibodies, 18 of them were selected for further studies. They bound potently and specifically to the SARS-CoV-2 RBD with subnanomolar to nanomolar affinities as measured by bio-layer interferometry (BLI) and ELISA (Fig. 2 and Supplementary Fig. 1). Most of the antibodies displayed the fast-on/slow-off kinetic pattern, except for n3063 which had the slow-on/slow-off binding kinetics with the slowest rate constant of association (*k*_on_ = 9.0×10^3^ M^-1^s^-1^) and dissociation (*k*_off_ = 4.5×10^−4^ s^-1^). The antibody n3021, in contrast, had the fastest association rate (*k*_on_ = 8.0×10^5^ M^-1^s^-1^), resulting in the highest binding affinity (K_D_ = 0.6 nM) among all tested antibodies.

**Figure 2.**
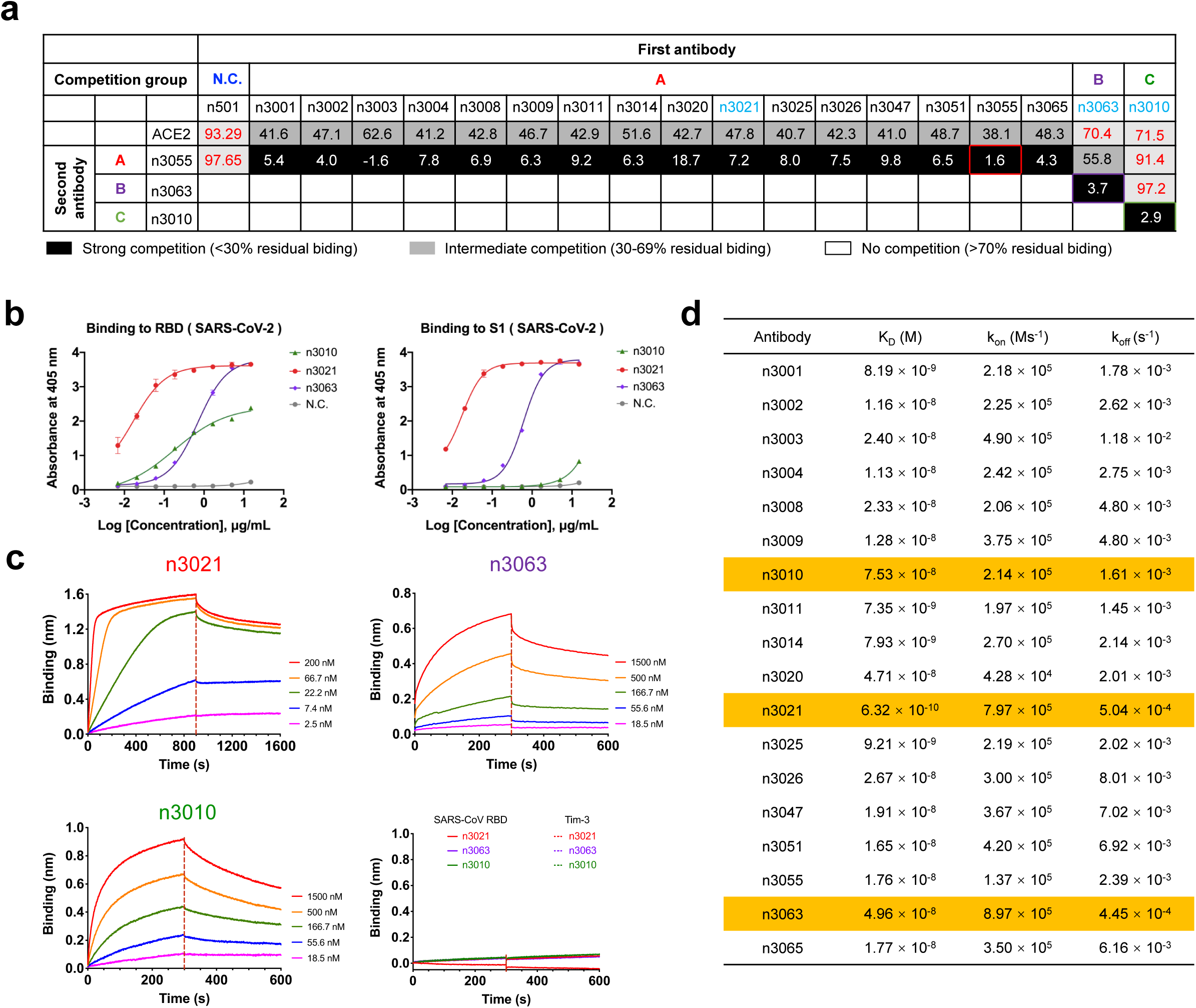
Characterization of human single-domain antibodies identified from antibody library using SARS-CoV-2 RBD as panning antigen. **a**, We tested 18 human single-domain antibodies targeting SARS-CoV-2 RBD in competition binding assays. Top: competition of human single-domain antibodies with ACE2 for RBD binding. The single-domain antibodies are displayed in three groups (A, B or C) based on a competition binding assay. The values are the percentage of binding that occurred during competition compared to non-competed binding, which was normalized to 100%, and the range of competition is indicated by the box colours. Black filled boxes indicate strongly competing pairs (residual binding <30%), grey filled boxes indicate intermediate competition (residual binding 30–69%), and white filled boxes indicate non-competing pairs (residual binding ≥70%). **b**, Binding of human single-domain antibodies to SARS-CoV-2 RBD or S1 as represented by competition group A antibody n3021, group B antibody n3063 and group C antibody n3010. **c**, Binding kinetics of competition groups A, B and C antibodies to SARS-CoV-2 RBD and binding specificity, as measured by BLI. **d**, List of binding properties of human single-domain antibodies. Association-rate (*k*_on_), dissociation-rate (*k*_off_) and affinity (K_D_) are shown. The representative single-domain antibodies of three groups are shown in yellow box.

To test whether these single-domain antibodies recognize different epitopes on RBD, the competition binding assays were performed, and the percentage of binding during competition compared to non-competed binding was quantitatively measured (Fig. 2a and Supplementary Fig. 2). We found that 18 antibodies could be divided into three competition groups (group A, B or C) that did not show any competition with each other. Most of the group A antibodies competed strongly with each other for binding to RBD, indicating that they recognized the same epitope. These results suggest that group A, B and C antibodies bound to different epitopes on RBD.

To further elucidate their binding epitopes, we measured the competition of single-domain antibodies and human ACE2 for binding to SARS-CoV-2 RBD (Fig. 2a and Supplementary Fig. 3). The antibodies n3063 (group B) and n3010 (group C) did not show any competition, while all the group A antibodies showed moderate competition with ACE2 for the binding to RBD. Therefore, the epitopes targeted by group A antibodies may be located within or adjacent to the ACE2-binding motifs of RBD.

**Figure 3.**
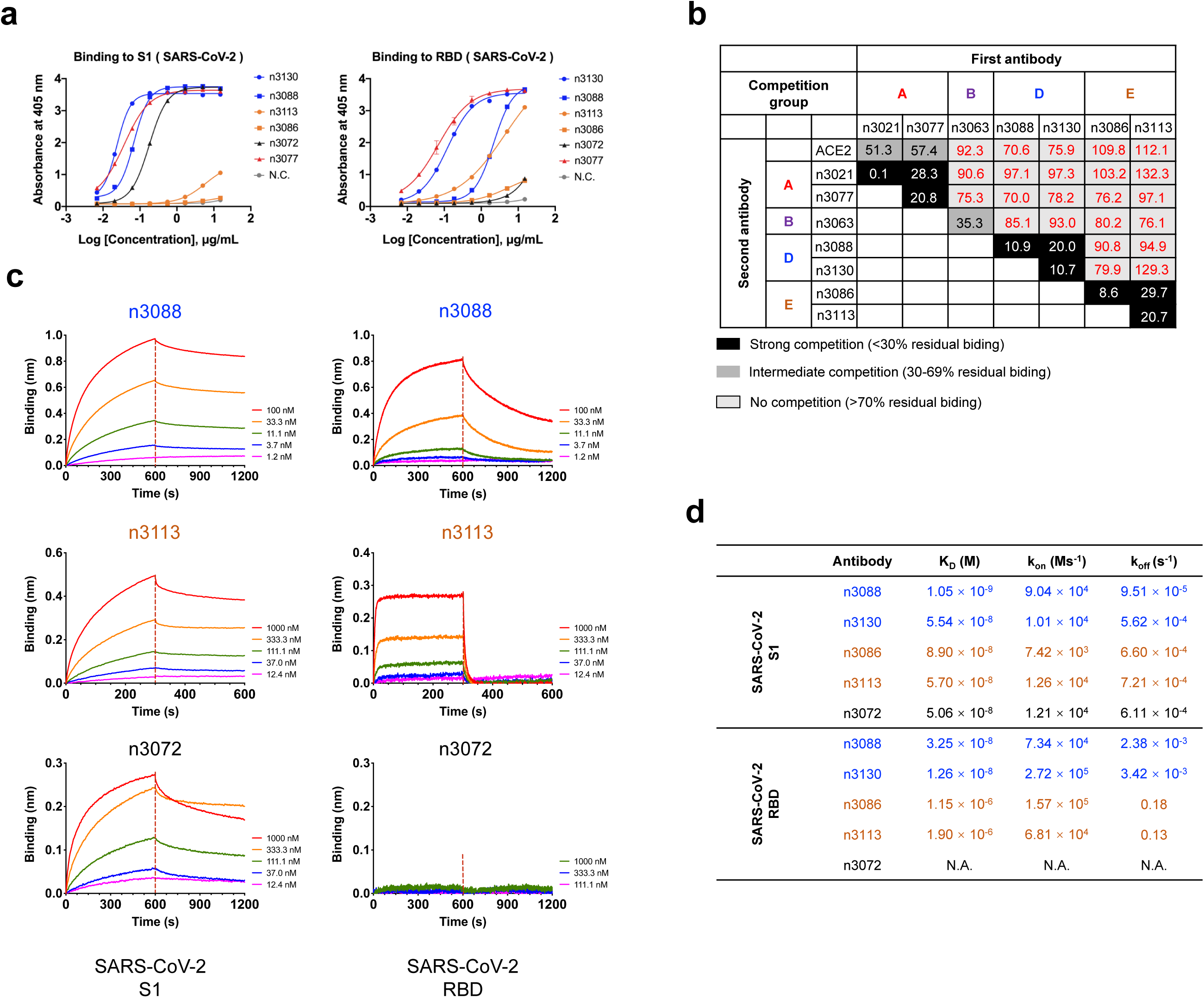
Characterization of human single-domain antibodies identified from antibody library using SARS-CoV-2 S1 as panning antigen. **a**, Binding capacities of single-domain antibodies to SARS-CoV-2 S1 or RBD measured by ELISA. **b**, Top: competition of human single-domain antibodies with ACE2 for RBD binding. The single-domain antibodies in another two competition groups (D or E) distinct from groups A, B, and C are displayed. **c**, The binding kinetics of competition groups D and E antibodies to SARS-CoV-2 S1 or RBD. **d**, List of binding properties of human single-domain antibodies. Association-rate (*k*_on_), dissociation-rate (*k*_off_) and affinity (K_D_) are shown. The group D is shown in blue, and group E is shown in orange.

To investigate the potential of these single-domain antibodies in neutralizing SARS-CoV-2, we measured their inhibitory activities in a well-established SARS-CoV-2 pseudovirus infection assay. To our surprise, none of these antibodies showed efficient neutralization at 50 μg/ml (data not shown), implying that moderate competition with ACE2 is not sufficient for potent SARS-CoV-2 neutralization. Interestingly, we also found that the group C antibody n3010 bound potently to SARS-CoV-2 RBD but did not show any binding to S1 protein, indicating that it recognized a cryptic epitope hidden in S1. These results taken together suggest that some non-neutralizing epitopes are relatively immunogenic in the isolated SARS-CoV-2 RBD, in contrast to that of SARS-CoV and MERS-CoV in which the neutralizing subregion were found to be highly immunogenic ^22^.

Next, we performed another set of panning using SARS-CoV-2 S1 protein instead of RBD as the target antigen in order to isolate single-domain antibodies targeting more diverse epitopes. A panel of 41 unique antibodies were identified after 4 rounds of panning. Notably, two of them were found to be identical to the previously isolated group A antibody n3021 or group B antibody n3063. The binding of 6 representative antibodies (n3072, n3077, n3086, n3088, n3113 and n3130) to SARS-CoV-2 S1 or RBD were measured by ELISA and BLI (Fig. 3 and Supplementary Fig. 4). Most of them showed potent binding to both S1 and RBD, while only one antibody, n3072, had strong binding to S1 but no binding to RBD (Fig. 3a,c). The competition binding assay suggests that n3077 recognized the same epitope as the previously identified group A antibodies (Fig. 3b and Supplementary Fig. 5). The other 4 antibodies could be divided into two distinct competition groups, group D (n3088, n3130) and group E (n3086, n3113). These two groups had no competition with each other or with previously identified antibodies for RBD binding, indicating that two novel epitopes on SARS-CoV-2 RBD were identified by this new panel of single-domain antibodies.

**Figure 4.**
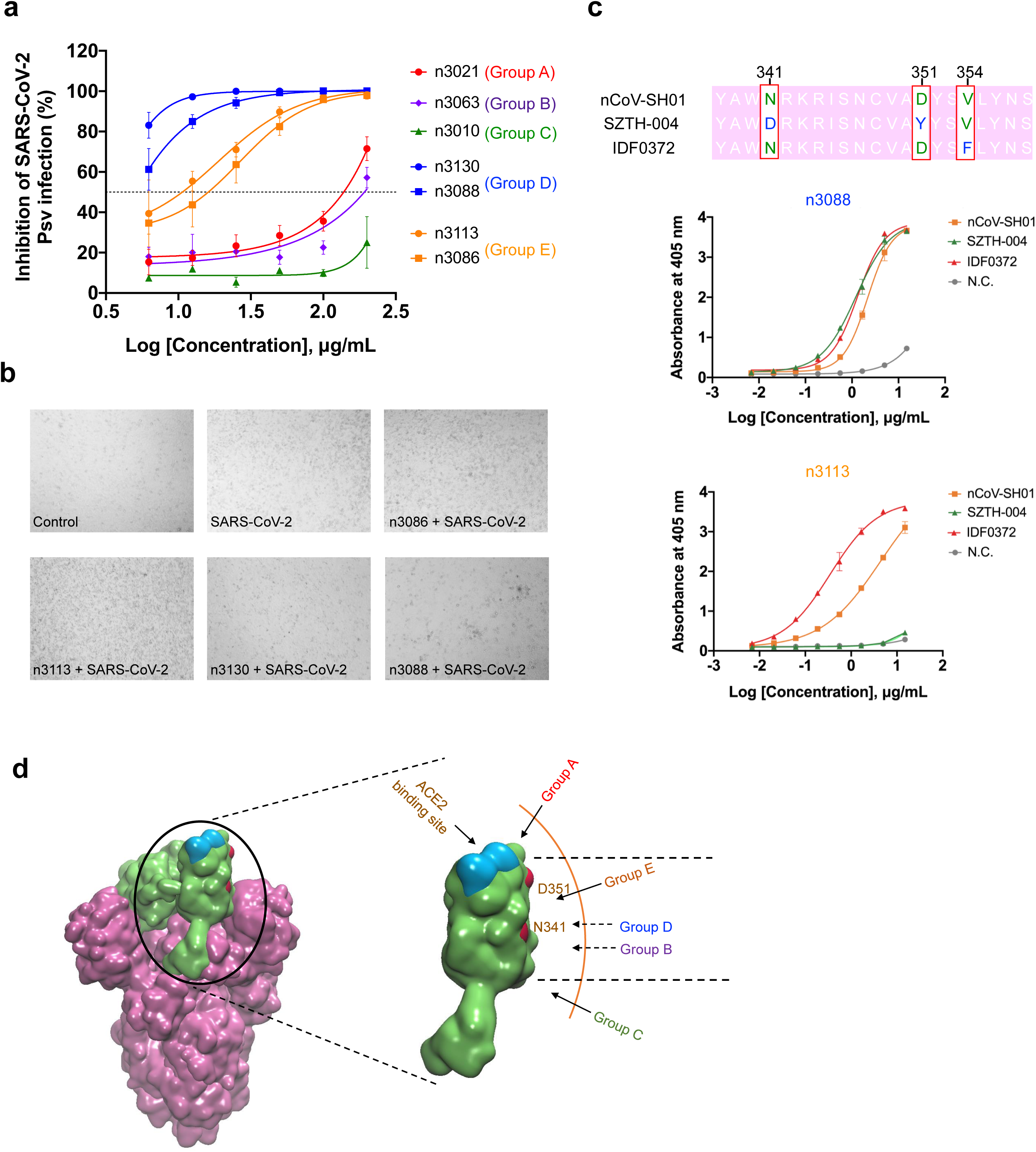
Neutralization activities of anti-SARS-CoV-2 human single-domain antibodies. **a**, Antibody-mediated neutralization against luciferase-encoding pseudotyped virus with spike protein of SARS-CoV-2. Pseudotyped viruses pre-incubated with antibodies at indicated concentrations were used to infect Huh-7 cells and inhibitory rates (%) of infection were calculated by luciferase activities in cell lysates. Dotted lines indicate inhibitory concentration at 50%. Error bars indicate mean ± s.d. from three independent experiments. **b**, The SARS-CoV-2 clinical isolate nCoV-SH01 was incubated with 20 μg/mL of single-domain antibodies for 1 h at 37°C prior to infection of Vero E6 cells. Subsequently, cytopathic effects (CPE) were observed daily and recorded on Day 3 post-exposure. **c**, Sequence alignment of three SARS-CoV-2 clinical isolates (nCoV-SH01, SZTH-004 and IDF0372) in which the mutations are hightlighted in red box, and binding capacity of neutralizing single-domain antibodies (group D antibody n3088 and group E antibody n3113) to RBD of three SARS-CoV-2 clinical isolates measured by ELISA, with an irrelevant protein (Tim-3) as control. **d**, Potential epitopes of antibodies from five competition groups A, B, C, D and E on RBD. RBD in the S protein of SARS-CoV-2 is shown green, and ACE2 binding site is colored blue. The two mutation sites (D351 and N341) of isolate SZTH-004 are shown in red.

We further measured the neutralization activities of these antibodies using the pseudovirus neutralizing assay. As shown in Fig. 4a, the group D antibodies exhibited potent neutralization of SARS-CoV-2 pseudovirus. The most potent antibody, n3130, could neutralize SARS-CoV-2 pseudovirus infection with >90% neutralization at 10 μg/ml. The other group D antibody n3088 neutralized ∼80% pseudovirus at 10 μg/ml. The group E antibodies n3086 and n3113 showed moderate neutralization activities, which inhibited SARS-CoV-2 pseudovirus infection in a dose-dependent manner with IC_50_ values of 26.6 and 18.9□µg/ml, respectively. The group A antibody n3021 and group B antibody n3063 could neutralize pseudovirus only at concentrations higher than 50 μg/ml, and group C antibody n3010 did not show evident neutralization activity. We next tested the neutralization of group D and E antibodies against live SARS-CoV-2 virus (Fig. 4b). Single-domain antibodies at 20 µg/ml were mixed with 200 PFU SARS-CoV-2 and observed for cytopathic effects (CPE) on Vero E6 cells. Similarly, no CPE was observed for n3130 and only very slight sign of CPE was found for n3088, while a significant level of CPE was detected in the wells containing group E antibodies.

It is very intriguing that the panning using SARS-CoV-2 S1 or RBD protein as antigen resulted in substantially different spectra of antibodies. The single-domain antibodies identified from S1 panning were very diverse, covering four distinct epitopes on SARS-CoV RBD (competition groups A, B, D and E). In contrast, most of the antibodies from RBD panning belonged to the competition group A, represented by n3021 which was also the most dominant clone after two rounds of panning. Furthermore, the group A antibodies showed moderate competition with ACE2 for RBD binding but insufficient to provide effective viral neutralization. This phenomenon is quite different from that of SARS-CoV, in which the dominance of an antigenic loop within RBD makes it relatively easy to isolate potent SARS-CoV neutralizing antibodies independent of repertoire, species, quaternary structure, and the technology used to derive the antibodies ^22^. Similarly, we previously used MERS-CoV S1 or RBD to isolate antibodies from a naïve antibody library, and the panning using either of the two antigens led to dominant enrichment of m336 and m336-like monoclonal antibodies, which precisely targeted 90% of the receptor binding site within RBD and neutralized the virus potently ^20^. It was proposed that viruses like SARS-CoV perhaps did not have sufficient evolutionary time to evolve their membrane glycoproteins to avoid direct immune recognition of a single site critical to the virus pathogenesis ^22^. It is noteworthy to point out that the difference in the immunogenicity of RBD was observed solely based on in vitro experiments, and it may not correlate with humoral immune responses in vivo. In this regard, it is imperative to investigate the immunogenic characteristics of SARS-CoV-2 RBD with special attention to the potentially antigenic and non-neutralizing epitopes. Besides, another interesting finding is that the SARS-CoV-2-specific neutralizing antibodies from competition groups D and E are not capable of competing with ACE2 for SARS-CoV-2 RBD binding (Fig. 3 and Supplementary Fig. 6). We found that the group E antibody n3113 did not exhibit any binding to the RBD of SARS-CoV-2 isolate SZTH-004 that had two mutations (N341D/D351Y) to the most prevelant isolate (Fig. 4c), indicating that the epitope of group D antibodies was located at a region surrounding N341 or D351 which is distinct from the ACE2 binding site (Fig. 4d). This phenomenon was also not observed in SARS-CoV. All the SARS-CoV-specific human neutralizing monoclonal antibodies, as far as we know, competed with ACE2 for binding to the spike protein. CR3022, a cross-reactive human monoclonal antibody that could neutralize SARS-CoV and was found to bind potently to SARS-CoV-2 RBD ^23^ but not capable of neutralizing SARS-CoV-2 ^24^. These findings confirmed the unique immunogenic profile of SARS-CoV-2. Further investigations are needed to understand the underlying mechanisms that govern these diverse sets of neutralizing and non-neutralizing SARS-CoV-2 antibodies, which may have important implications for the development of effective vaccines.

The fully human single-domain antibodies offer the potential for prevention and treatment of COVID-19. First, antibodies derived entirely of human sequences would be less immunogenic than camelid or humanized nanobodies, leading to improved safety and efficacy when used in humans. Indeed, despite humanization, caplacizumab, the first nanobody approved by FDA still contains multiple camelid residues to maintain the antigen binding affinity. Second, the small size and favorable biophysical properties allows for large-scale production of single-domain antibodies within a few weeks in prokaryotic expression systems, and thus enables rapid implementation in an outbreak setting. Furthermore, single-domain antibodies could be delivered to the lung via inhalation, which may offer considerable advantages for treatment of COVID-19 including fast onset of action, low systemic exposure, and high concentration of therapeutics at the site of disease. Lastly, single-domain antibodies can be used alone or synergistically with other neutralizing antibodies. Their small size making them ideal building block for generation of bispecific or multi-specific antibodies to prevent the appearance of viral escape mutants. They can also be easily engineered to further increase the neutralization activity by increasing binding moieties. For instance, the trivalent nanobody ALX-0171 was found to have 6,000-fold increased neutralization potency against RSV-A and >10,000-fold against RSV-B compared to its monovalent format ^25^.

In summary, we report here the development of a versatile platform for rapid isolation of fully human single-domain antibodies, and its application for screening of antibodies against SARS-CoV-2. A variety of single-domain antibodies have been isolated targeting five types of epitopes on SARS-CoV-2, and the antibody n3130 was found to potently neutralize both pseudotyped and live virus. These antibodies may represent promising candidates for prophylaxis and therapy of COVID-19, and also serve as reagents to facilitate the vaccine development.

## Author contributions

TY, YW and CT conceived and designed the study. YW, TY and CL performed most of the experiments with assistance from XT, ZW, and YK. SX, LL and SJ performed pseudovirus neutralization assay. CG, RZ and YX performed live SARS-CoV-2 neutralization assay. TY and YW integrated the data and wrote the manuscript. All authors reviewed and approved the final version of the manuscript.

## Acknowledgments

We thank Chengfeng Qin from Beijing Institute of Microbiology and Epidemiology, Zhenlin Yang, Ailing Huang and Shanshan Zhou from our group, Yang Wu and Yuyan Wang from BSL-3 laboratory of Fudan University, and the staff from Core Facility of Microbiology and Parasitology, Shanghai Medical College, Fudan University, for the help with experiments. This work was supported by grants from the National Key R&D Program of China (2019YFA0904400), National Natural Science Foundation of China (81822027, 81630090), National Megaprojects of China for Major Infectious Diseases (2018ZX10301403), and Chinese Academy of Medical Sciences (2019PT350002).

## Declaration of interest statement

No potential conflict of interest was reported by the authors.

## Figure legend

**Figure S1.**
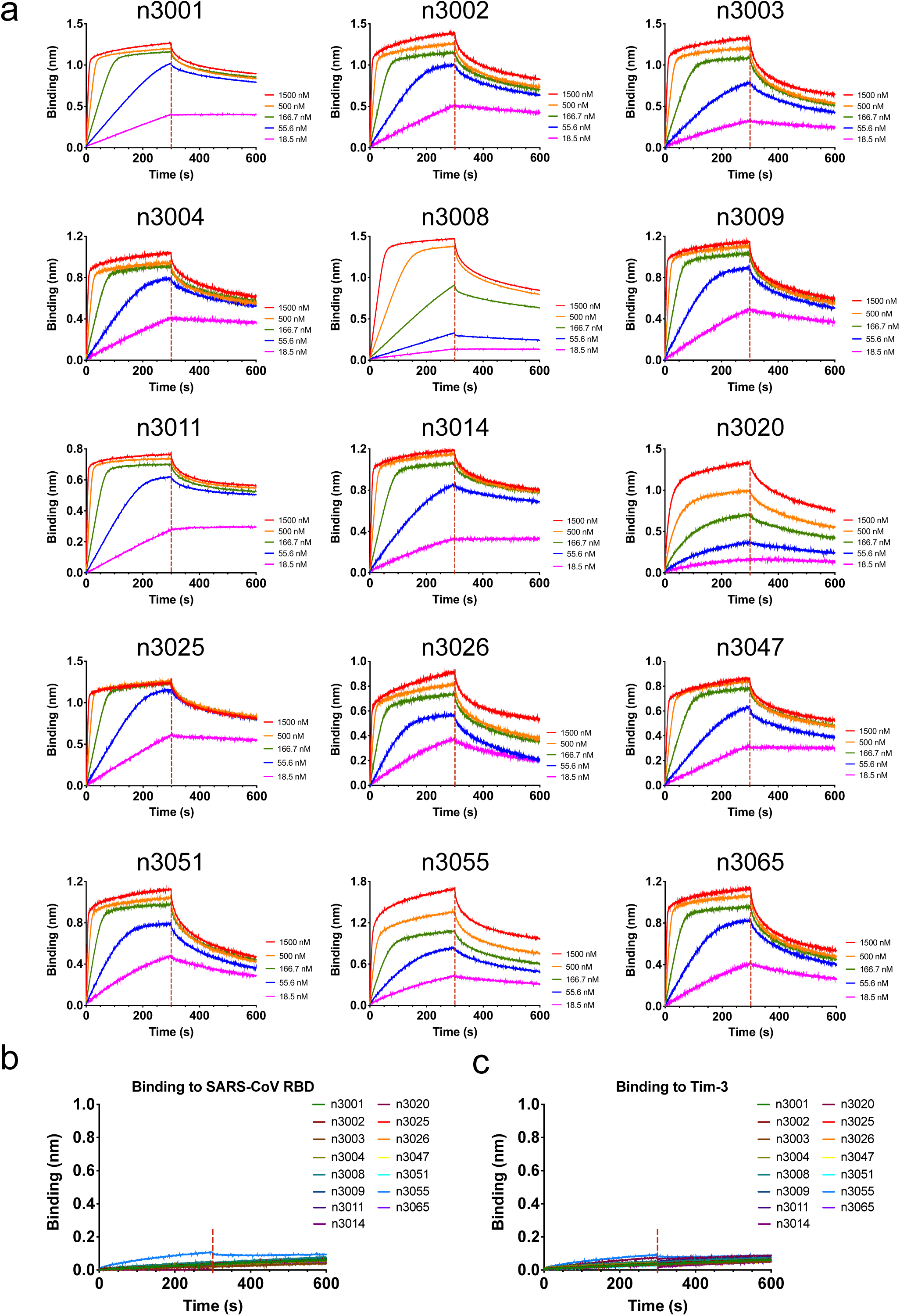
Binding kinetics of 15 human single-domain antibodies to SARS-CoV-2 RBD (a), SARS-CoV RBD (b), or control antigen (c), as measured by BLI using OctetRED96. Biotinylated SARS-CoV-2 RBD, SARS-CoV RBD or control antigen (Tim-3) was immobilized on SA biosensors. The analytes consisted of serial dilutions of single-domain antibodies between 22.5 μg/mL and 0.3 μg/mL or a single concentration at 15 μg/mL. Binding kinetics were evaluated using a 1:1 Langmuir binding model by Fortebio Data Analysis 10.0 software.

**Figure S2.**
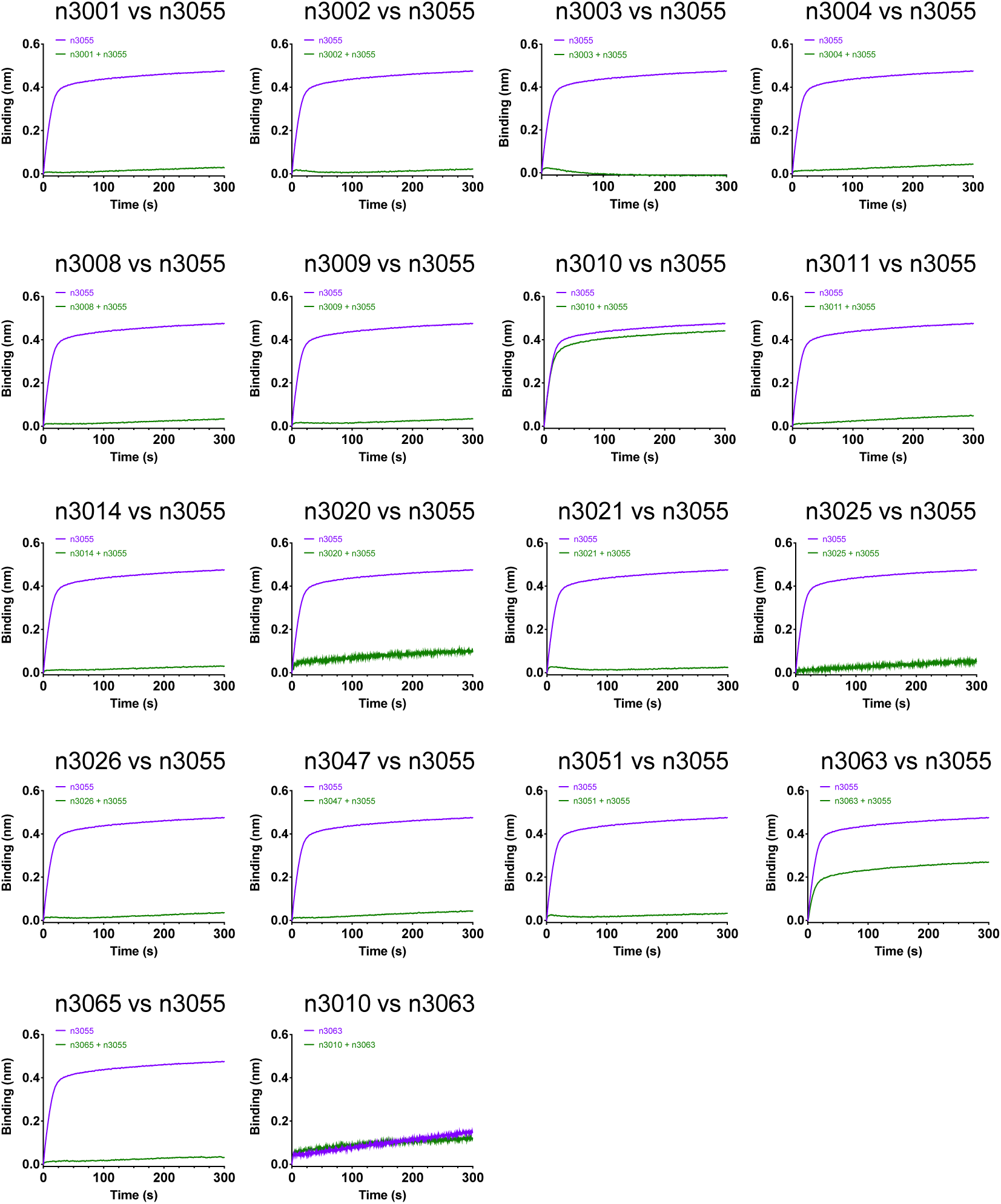
Competition of 18 human single-domain antibodies identified from antibody library using SARS-CoV-2 RBD as panning antigen, as measured by BLI. The competition assay was performed among 18 human single-domain antibodies for binding to RBD. Immobilized SARS-CoV-2 RBD was first saturated with 15 μg/mL of the first testing antibody. The capacity of the second antibody binding to RBD was monitored by measuring further shifts after injecting the second single-domain antibody (15 μg/mL) in the presence of the first single-domain antibody (15 μg/mL). The grams show binding patterns of the second single-domain antibody to SARS-CoV-2 RBD with (green curve) or without (purple curve) prior incubation with each testing single-domain antibody.

**Figure S3.**
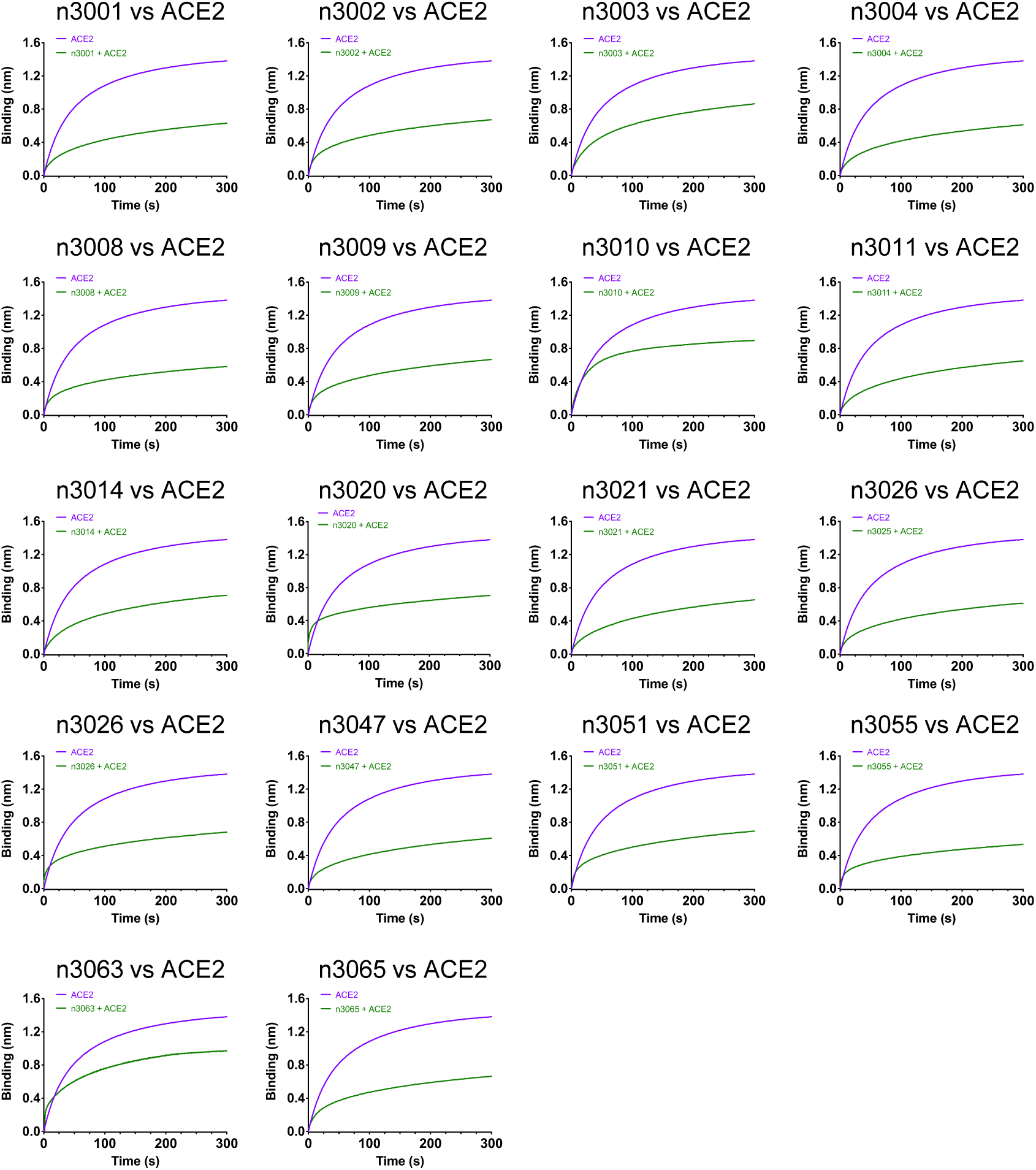
Human single-domain antibodies and ACE2 competition for binding to SARS-CoV-2 RBD. Immobilized SARS-CoV-2 RBD was first saturated with 15 μg/mL of the testing single-domain antibodies. The capacity of ACE2 binding to RBD was monitored by measuring further shifts after injecting the ACE2 (17 μg/mL) in the presence of the testing single-domain antibody (15 μg/mL). The grams show binding patterns of ACE2 to SARS-CoV-2 RBD with (green curve) or without (purple curve) prior incubation with each testing single-domain antibody.

**Figure S4.**
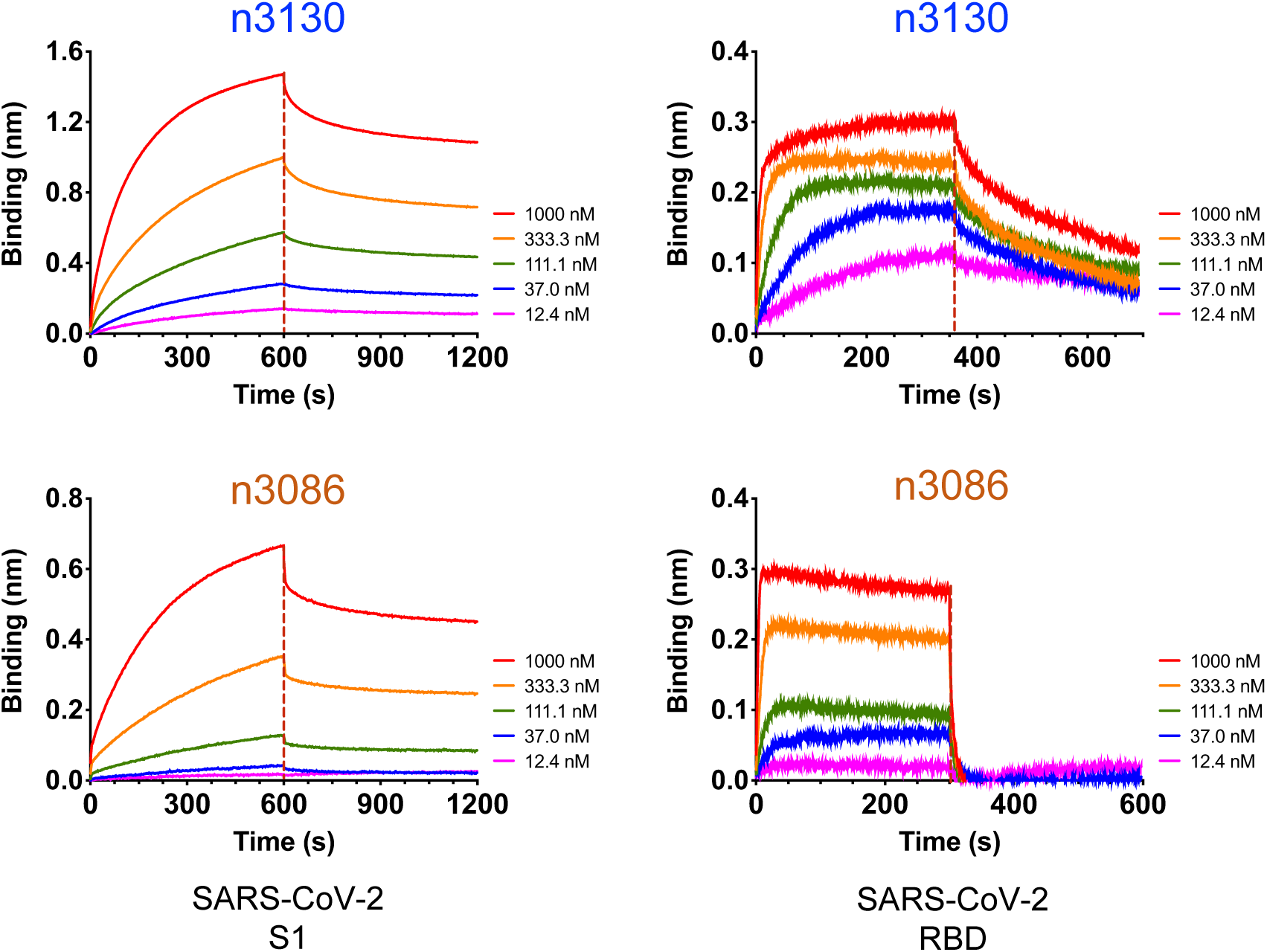
Binding kinetics of n3130 and n3086 to SARS-CoV-2 S1 and RBD. Sensors immobilized SARS-CoV-2 S1 or RBD were incubated with five dilutions of n3130 and n3086 for 300 s or 600 s, and then transferred into kinetic buffer for dissociation.

**Figure S5.**
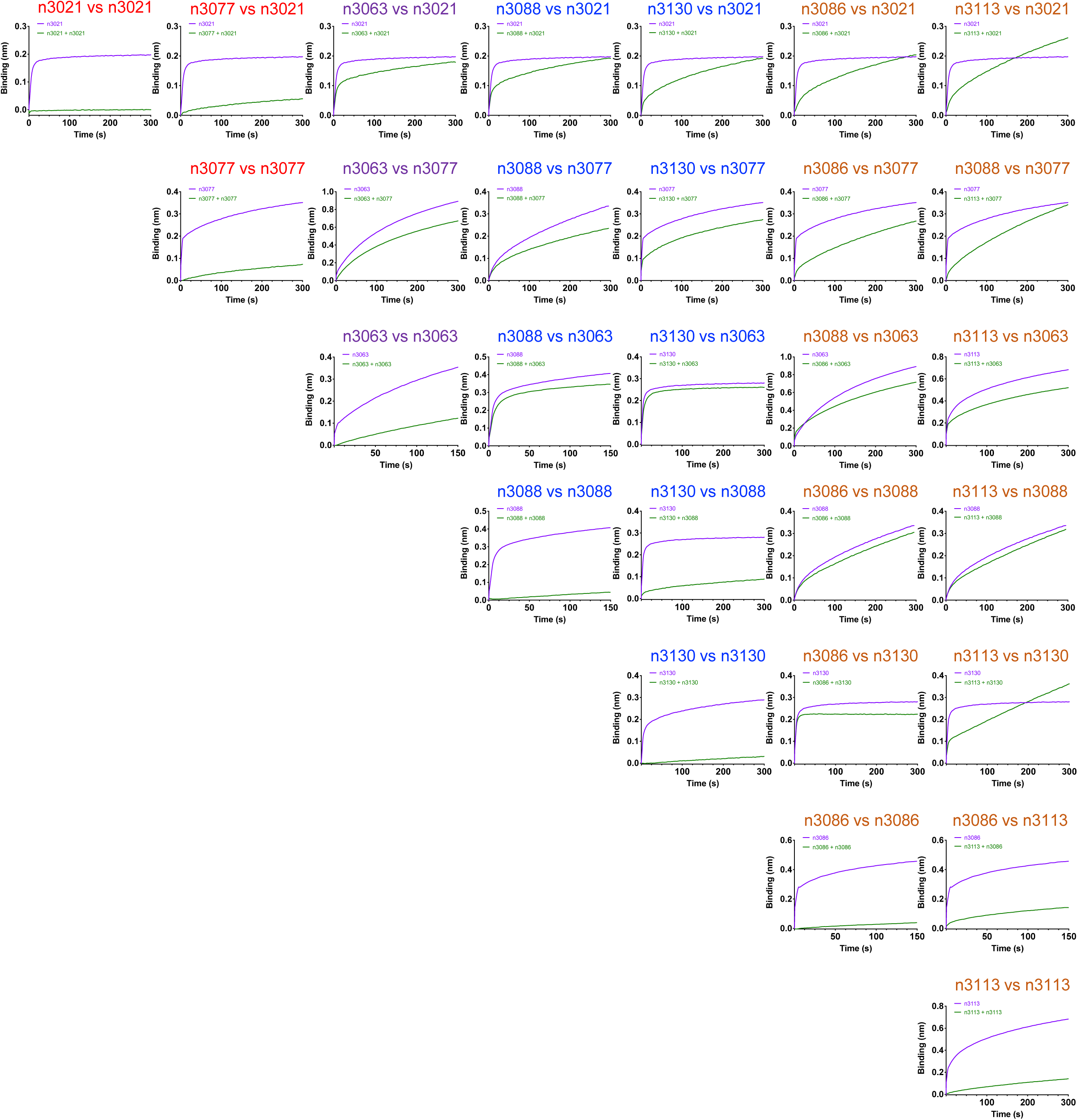
Competition of human single-domain antibodies identified from antibody library using SARS-CoV-2 S1 as panning antigen, as described in legend of Fig S2.

**Figure S6.**
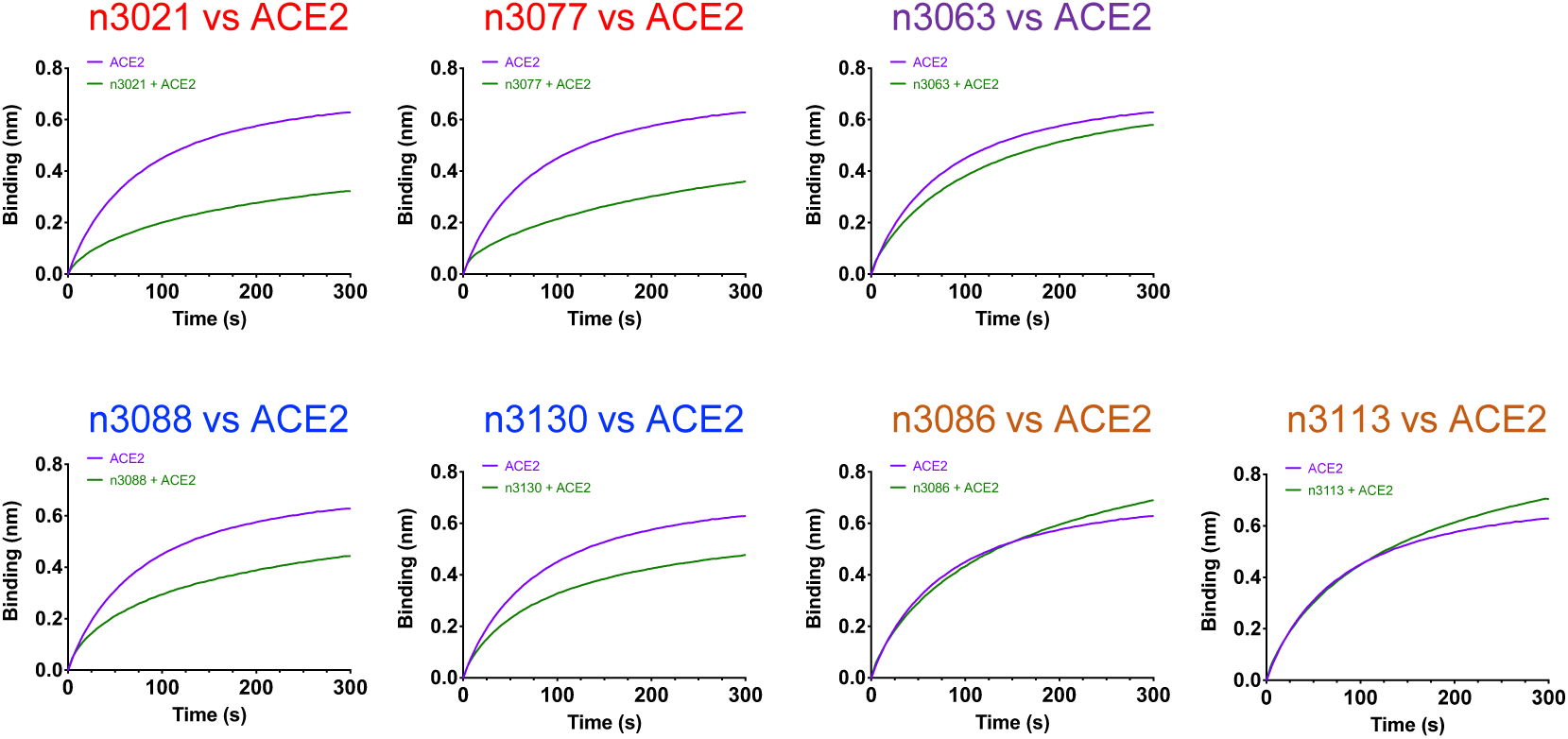
Human single-domain antibodies of group A, B, D or E and ACE2 competition for binding to SARS-CoV-2 RBD, as described in legend of Fig S3.

