## Supplementary material for "Fully human single-domain antibodies against SARS-CoV-2": Methods

**Protein expression and purification.** The recombinant receptor binding domain (RBD) sequences encoding amino acids R319-F541 of SARS-CoV-2 (isolate nCoV-SH from GISAID accession no. EPI_ISL_402124, isolate SZTH-004 from GISAID accession no. EPI_ISL_406595, isolate IDF0372 from GISAID accession no. EPI_ISL_406596/97) , amino acids R306-F527 of SARS-CoV (isolate:WH20) were cloned into pSecTag2B expression vector with an additional C-terminal Fc fragment of human IgG1 and AviTag in tandem and transiently expressed in Expi293 cells as a secreted protein. After 3 days, supernatants were harvested by centrifuging the culture at 2500 g for 15 min and filtering the supernatant with a 0.22 mm vacuum filter. The protein was purified using protein G resin (GE Healthcare). Equilibration and wash steps were performed with phosphate buffered saline solution (PBS) (Hyclone) and protein were eluted in 0.1 M glycine pH 2.7. The eluates were pH equilibrated to 7.4 using 1.0 M Tris-HCl pH 9.0 and immediately buffer-exchanged into PBS and concentrated using an Amicon ultra centrifugal concentrator (Millipore) with a molecular weight cut-off of 10 kDa. Purity was estimated as 95% by SDS–polyacrylamide gel electrophoresis, and protein concentration was measured using the NanoDrop 2000 spectrophotometer (Thermo Fisher). The biotinylated proteins used in bio-panning and BLI experiment were prepared by the BirA biotin-protein ligase in PBS for 30 min at 30 °C according to the manufacturer’s instructions (Avidity), which adds biotin covalently to AviTag in a highly specific manner, or by non-specific labeling using sulfo-NHS-LC-Biotin (Thermo Fisher). The human single-domain antibody (single-domain antibody) and camelid nanobody were subcloned into the pComb3x vector with C-terminal tags comprising a hexahistidine (His6) tag and a Flag tag in tandem. Their expression was performed in *Escherichia coli* HB2151 bacterial culture at 30°C for 14 h accompanied with 1 mM IPTG. The cells were harvested and lysed by Polymyxin B (Sigma-Aldrich) at 30°C for 0.5 h. Supernatant was obtained by centrifugation at 8000 rpm for 10 min and loaded over Ni-NTA as manual described (GE Healthcare). Resin was washed by washing buffer [10 mM Na_2_HPO_4_, 10 mM NaH_2_PO_4_ (pH 7.4), 500 mM NaCl and 20 mM imidazole] and protein were eluted in elution buffer [10 mM Na_2_HPO_4_, 10 mM NaH_2_PO_4_ (pH 7.4), 500 mM NaCl and 250 mM imidazole]. The collected pure fractions were immediately buffer-exchanged into PBS and concentrated using an Amicon ultra centrifugal concentrator (Millipore) with a molecular weight cut-off of 3 kDa.

**Determination of melting and aggregation temperature.** Thermal stability and colloidal stability were assessed by using a Uncle/UNit system (Unchained Labs, Pleasanton, CA). Briefly, the static light scattering (SLS) at 473 nm was used as an indicator for “colloidal stability”, reporting the onset of aggregation temperature (Tagg), which can be defined as the temperature at which the measured scatter reaches a threshold that is approximately 10% of its maximum value. The changes in the SLS signal represented changes in the weight average molecular mass observed due to protein aggregation. Thermal stability was evaluated at an intrinsic fluorescence intensity ratio (350/330 nm) by measuring the temperature of the on-set of melting. The samples at a concentration of 0.5 mg/mL were heated from 20 °C to 95 °C using 1 °C increments, with an equilibration time of 60 s before each measurement. Measurements were made in duplicates.

**Construction of human domain antibody library and bio-panning.** For the construction of the large and highly diverse full-human single-domain antibody library, grafting of the CDR regions of heavy chain from several naïve antibody libraries was carried out by PCR using specific oligonucleotides and combined with framework sequences from the germline IGHV3-66*01 subfamily by a series of overlap extension PCR (OE-PCR). The final PCR products were cloned into phagemid pComb3x allowing the expression of C-terminal His6-Flag tagged VHs. The recombinant vector was electro-transformed into competent TG1 bacteria and grown in 2 × YT medium with 100 μg/mL ampicillin and 2% (w/v) glucose. When the optical density at 600 nm (OD_600_) reached 0.6, TG1 were infected with M13KO7 helper phages for 1 h at 37°C. The TG1 were harvested and resuspended in 2 × YT medium with ampicillin and kanamycin (100 μg/mL), and the bacteria were cultured overnight at 30°C. Next day, the cultures were centrifuged and phages were precipitated from the supernatant by adding 5% (w/v) polyethylene glycol (PEG) 8000-NaCl (20 % PEG8000, 2.5 mol/L NaCl). Precipitated phages were collected by centrifugation, resuspended in sterile PBS, and frozen in aliquots of 10^13^ CFU at -80°C. Panning protocols were carried essentially out as described previously^1,2^. For panning of the library against target molecules, 5 µg recombinant biotinylated antigen was utilized in round 1, 4 µg was utilized in round 2, 2 µg was utilized in round 3, and 1 µg was utilized in round 4, which were immobilized on streptavidin-coated magnetic beads (Invitrogen). About 1×10^12^ phage particles were used for each bio-panning. Positive clones expressing human single-domain antibody were identified from the richen rounds of panning by using monoclonal ELISA.

**Enzyme-linked immunosorbent assay (ELISA).** Costar half-area high binding assay plates (Corning #3690) were coated with purified protein at 100 ng/well in PBS overnight at 4°C and blocked with PBS buffer containing 3% milk powder (w/v) at 37°C. Serially diluted antibody solutions were added and incubated for 1.5 h at 37°C. The bound antibodies were detected with monoclonal anti-flag-HRP antibody (Sigma-Aldrich). The enzyme activity was measured with the subsequent addition of substrate ABTS (Invitrogen) and signal reading was carried out at 405 nm using a Microplate Spectrophotometer (Biotek).

**Biolayer interferometry (BLI) binding assays.** All BLI was carried out on an OctetRED96 device (Pall FortéBio) and assays were carried out in 96-well format in black plates (Greiner). For the binding kinetics of single-domain antibodies with SARS-CoV-2 S1, the S1 protein at 15 μg/mL buffered in sodium acetate (pH 5.0) was immobilized onto activated AR2G biosensors (Pall FortéBio) and incubated with threefold serial dilutions of single-domain antibodies in kinetics buffer (PBS buffer supplemented with 0.02% Tween 20). The experiments included the following steps at 37 °C: (1) equilibration (60 s); (2) activation of AR2G by 1-ethyl-3-(3-dimethylaminopropyl)carbodiimide hydrochloride/N-hydroxysucci-nimide (300 s); (3) immobilization of S1 protein onto sensors (100 s); (4) quenching with ethanolamine (300 s); (5) baseline in kinetics buffer (120 s); (6) association of antibodies for measurement of *k*_on_ (300-600 s); and (7) dissociation of antibodies for measurement of *k*_off_ (300-600 s). The curves were fitted by a 1:1 binding model using FortéBio Data Analysis software 10.0. For measuring binding kinetics of single-domain antibodies with SARS-CoV-2 RBD, Avi-tagged recombinant RBD was biotinylated with the BirA biotinylation kit (Avidity), diluted in kinetics buffer and immobilized on streptavidin (SA) coated biosensors (Pall FortéBio) at ~50% of the sensor maximum binding capacity. Baseline was established in kinetics buffer and loaded biosensors were dipped into wells containing serial dilutions of single-domain antibodies for 300-600 s. RBD:single-domain antibody complexes were then allowed to dissociate in kinetics buffer. After reference subtraction, apparent binding kinetic constants were determined by fitting the curves to a 1:1 binding model using the Data Analysis software 10.0 (FortéBio). Real-time interactions between purified SARS-CoV RBD or an irrelevant protein (Tim-3) as control and single-domain antibodies were determined. The streptavidin-coated biosensors were loaded biotinylated SARS-CoV RBD or Tim-3, and then incubated with 15 μg/mL of single-domain antibodies in kinetics buffer and binding was measured after 300 s of association.

**Binding competition assays.** The epitopes of anti-SARS-CoV-2 single-domain antibodies were initially mapped by binding competition assays using biolayer interferometry (BLI). Sensor tips loaded with SARS-CoV-2 RBD, as described above, were immersed into wells containing the first competing single-domain antibody at a concentration (15 μg/mL) necessary to reach binding saturation after 300 s. Next, biosensors were dipped into wells containing the second single-domain antibody with the same concentration at 15 μg/mL or 17 μg/mL of ACE2, in the presence of the first competing single-domain antibody, and binding was measured after 300 s of association. The signal obtained for binding of the second antibody in the presence of the first antibody was expressed as a percentage of the uncompeted binding of the second antibody that was derived independently. The antibodies were defined as competing if the presence of first antibody reduced the signal of the second antibody to less than 30% of its maximal binding capacity, and non-competing when binding was greater than 70%. A level of 30-69% was considered intermediate competition.

**Pseudotyped virus neutralization.** To determine the neutralization activity of single-domain antibodies, the pseudotyped virus neutralization assay was performed. Briefly, 293 T cells were co-transfected with expression vectors of pcDNA3.1-SARS-CoV-2-S (encoding SARS-CoV-2 S protein) and pNL4-3.luc.RE bearing the luciferase reporter-expressing HIV-1 backbone, as previously described^3^. The supernatants containing SARS-CoV-2 pseudotyped virus were harvested. Serial dilutions of single-domain antibodies in DMEM supplemented with 10% fetal calf serum were incubated with pseudoviruses at 37°C for 1 h and then the mixtures were added in monolayer Huh-7 cells (10^4^ per well in 96-well plates). Twelve hours after infection, culture medium was refreshed and then incubated for an additional 48 hours. The luciferase activity was calculated for the detection of relative light units using the Bright-Glo™ Luciferase Assay System (Promega) in Ultra 384 luminometer (Tecan). A nonlinear regression analysis was performed on the resulting curves using Prism (GraphPad) to calculate half-maximal inhibitory concentration (IC_50_) values.

**Live virus neutralization.** Viral cytopathic effect (CPE)-based neutralization assay was conducted to determine the neutralizing human single-domain antibodies against live SARS-CoV-2 virus. In brief, 1×10^4^ Vero E6 cells were seeded in 96-well plates and cultured for overnight at 37°C. Single-domain antibodies were diluted at a final concentration of 20 μg/mL in culture medium and mixed with 200 PFU of SARS-CoV-2 (isolate nCoV-SH01). Following incubation for 1 h at 37°C, the virus and antibody mixtures were added to the Vero E6 cells, and medium was replaced with fresh DMEM 2 h later. Viral cytopathic effect (CPE) was observed daily and recorded on Day 3 post-infection. The neutralization experiment was performed under BSL-3 conditions. A minimum of 3 independent experiments was performed.

**Statistical analysis.** Binding experiments are presented as the mean values ± s.d. calculated from two independent experiments. Neutralization is the geometric mean of the IC_50_ values calculated using four-parameter logistic regression from at least two independent experiments performed in triplicate. All data were graphed using Prism software (version 8, GraphPad Software).
